# A Snakemake-based bacterial whole genome comparison pipeline for multi-group clinical isolates

**DOI:** 10.64898/2026.06.25.734687

**Authors:** Hoeyoung Kim, Ho Seok Sim, Jaeeun Kim, Kwangsoo Kim, Jinki Yeom

## Abstract

Organisms have continuously evolved in response to environmental conditions. Pathogenic bacteria evolve under host and environmental pressures, reshaping their genomes through insertions, inversions, deletions, and duplications during infection. In clinical settings, phenotypic traits of pathogenic bacteria such as virulence or antimicrobial resistance directly affect disease severity, transmission, and treatment. Conventional genotyping provides insights into genomic relatedness but does not always align with these clinically relevant traits, limiting its utility for phenotype-driven interventions. Here, we develop ABComp (Assembly polishing and Bacterial whole-genome Comparison for multi-group clinical isolates), a modular and Snakemake-based workflow for phenotype-driven comparative genomics. ABComp automates assembly polishing, group-wise pangenome analysis, and enables flexible pathogenic marker discovery through user-defined comparisons. We validated ABComp using a *Klebsiella pneumoniae* ground truth dataset stratified by yersiniabactin presence and successfully recovered the entire locus as a group-specific core marker. By applying ABComp to another dataset of clinical isolates with experimentally measured virulence, we discovered the ferric citrate (Fec) uptake system as a potential marker specific to a hypervirulent group. These results demonstrate ABComp’s utility in uncovering phenotype-linked genomic markers with clinical significance, supporting targeted treatments and rapid diagnosis.

## Introduction

Bacterial infections remain one of the leading causes of mortality worldwide, driven by the emergence of multi-drug resistant (MDR) strains^1,2^. MDR bacteria, such as carbapenem-resistant *Enterobacterales* (CRE) and *Acinetobacter baumannii* (CRAB), have developed resistance to last-resort antibiotics through genomic evolution processes such as horizontal gene transfer, complicating treatment options^3–5^. In some cases, populations of MDR bacteria have evolved into hypervirulent strains that pose significant challenges due to their ability to evade host immune defenses and establish severe infections^6,7^. Although virulence is the most critical factor for clinical outcomes, it remains largely unknown how MDR bacteria evolve their virulence during infections and how we can detect critical virulence factors from them. Thus, genomic analysis of bacterial clinical isolates has become essential not only for unraveling the genetic basis of resistance but also for identifying virulence factors critical for pathogenesis and treatment strategies. In this study, we developed a genomic analysis tool to detect critical virulence factors from clinically isolated MDR bacteria.

Recent advancements in whole genome sequencing (WGS) have enabled a more extensive study of both antibiotic resistance and virulence mechanisms in bacteria. Especially, the development of long-read sequencing technologies, such as Oxford Nanopore Technologies^8^ (ONT) and Pacific Biosciences^9^ (PacBio), have provided more continuous genome assemblies, overcoming the limitations of short-read-only assemblies in resolving repetitive regions^10–12^. However, long-read platforms, particularly ONT long reads, are prone to base errors, exhibiting higher insertion and deletion rates especially in homopolymeric regions^13^. To mitigate this limitation, hybrid assembly methods, which combine long-read data with highly accurate short reads (e.g., Illumina), have become a standard to improve both genome continuity and basecall accuracy^14,15^. These high-quality genome assemblies enable accurate downstream analyses, such as gene annotation, pangenome analysis, and virulence factor screening.

Currently, clinically isolated bacteria are categorized into groups by molecular typing methods that provide information about bacterial community structure and outbreak tracking. One widely used molecular typing method is Multilocus Sequence Typing (MLST), which classifies isolates based on sequence variations across a set of housekeeping genes^16^. While this approach provides valuable insights into bacterial phylogeny, it does not always correlate with clinical phenotypes, such as virulence or antibiotic resistance^17^. Given this limitation, a more targeted approach is required in clinical settings to define groups based on clinically meaningful phenotypes. These may include high- and low-virulence strains based on patient survival data, drug-resistant versus susceptible isolates, or strains associated with distinct infection sites or pathogenesis mechanisms. Comparative genomic analysis of such phenotype-driven groups is critical for identifying group-specific virulence markers that may not be captured by conventional genotyping approaches.

A range of automated bacterial WGS pipelines now span genome assembly, annotation, and antimicrobial-resistance or virulence-gene screening, including frameworks such as TORMES^18^, Bactopia^19^, and rMAP^20^, and, more recently, long-read-oriented workflows such as StrainCascade^21^. These tools excel at per-isolate profiling and population-scale phylogenetics, and some can be extended to pangenome construction^22^ and, through dedicated pan-genome-wide association tools such as Scoary^23^ or Pyseer^24^, to statistical genotype–phenotype association. Other pipelines target specific tasks such as variant calling^25^ or core-genome annotation^26^, stopping at genotyping or tree building. Across these tools, however, phenotype-driven discovery remains limited. Statistical approaches such as pan-GWAS require large sample sizes and explicit population-structure correction, while most end-to-end pipelines offer no phenotype-linked marker discovery at all. As shown in Table 1, no existing framework integrates hybrid assembly polishing, group-stratified pangenome reconstruction, and deterministic between-group marker discovery in a single workflow (Table 1). ABComp was developed to occupy this niche: a Snakemake workflow that takes user-defined phenotypic groups and reports group-specific core markers through transparent presence/absence comparison with additional functional enrichments, complementing rather than replacing statistical association approaches.

In this study, we developed ABComp (Assembly polishing and Bacterial whole-genome Comparison for multi-group clinical isolates), a Snakemake^27^-based workflow that automates the process from genome assembly polishing to group-wise pangenome analysis and pathogenic marker discovery. Unlike conventional comparative genomics tools, ABComp enables user-defined group comparisons, allowing the discovery of potential genomic markers that can define clinically relevant phenotypes. The tool is modular and flexible, accepting both raw sequencing reads and pre-assembled genomes as initial input. By integrating these functionalities, ABComp facilitates reproducible, scalable, and clinically actionable genomic analyses, making it a powerful tool for both research and diagnostic applications. Therefore, ABComp provides insights into virulence evolution of MDR bacteria and offers new strategies for their control.

## Results

### ABComp is a modularized pipeline for phenotype-driven bacterial genomic analysis

Pathogenic bacteria continuously evolve their genomes to survive in hosts, which increases the severity of pathogenesis through hyperactivation of virulence and antibiotic resistance mechanisms. Specifically, antibiotic-resistant bacteria induce virulence factors that cause sepsis and promote death in nosocomial settings^28^. Recently, genomes of many antibiotic-resistant bacteria have been sequenced and deposited in public databases^29,30^. However, until now, we have not had common tools to analyze and identify virulence factors from antibiotic-resistant bacteria isolated from patients. In addition, the deposited raw sequencing data of whole genomes from antibiotic-resistant bacteria are very diverse, which creates a bottleneck for detecting common virulence factors and predicting evolution from clinically isolated bacteria.

To enable phenotype-driven comparative bacterial genomic analysis for detecting evolved virulence factors in clinically isolated bacteria, we developed the ABComp pipeline, which is modularized into three core components. The first two modules, Assembly Polishing and Comparative Genomics, are automated via Snakemake (Fig. 1a,b), while the third module, Pathogenic Marker Discovery, allows between-group comparison of user-selected groups to identify group-specific pathogenic markers (Fig. 2a,b).

**Figure 1.**
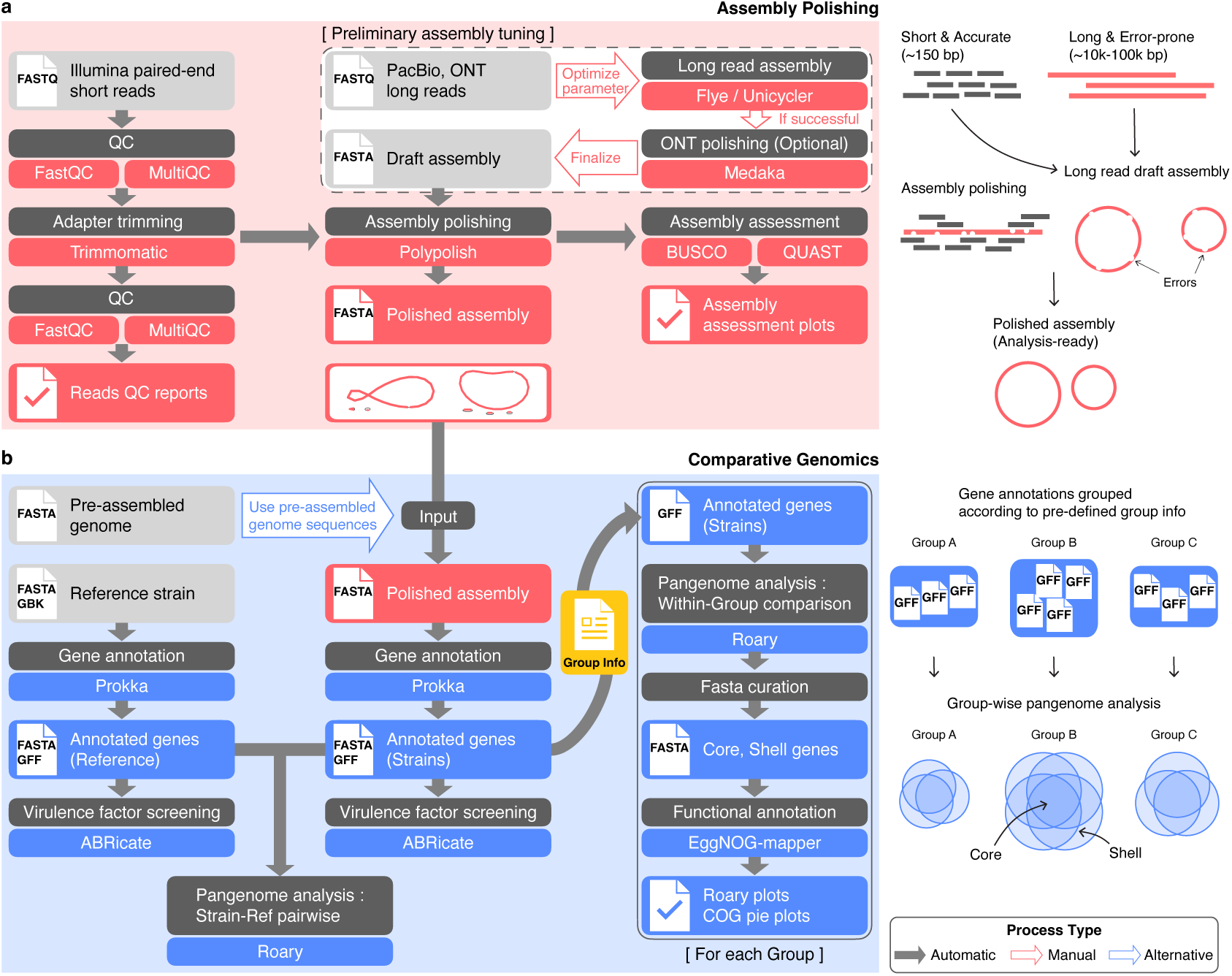
Overview of the ABComp workflow. ABComp consists of two main modules: **a** The Assembly Polishing module accepts short-read (Illumina) data and draft assembly generated from long-read sequencing technologies (e.g., Pacific Biosciences or Oxford Nanopore Technologies). Illumina reads undergo quality control, adapter trimming, are then used to polish the draft assembly to improve base-level accuracy. **b** The Comparative Genomics module begins with gene annotation and screening for virulence and resistance genes. The annotated genes are grouped based on a user-defined configuration file, and subjected to pangenome analysis to infer core and shell gene sets. Each gene set is curated as a sequence file, then undergoes functional annotation. Parallely, each strain is compared to the reference strain in a pairwise manner. Alternatively, users may provide pre-assembled genomes as input to begin directly from the module.

**Figure 2.**
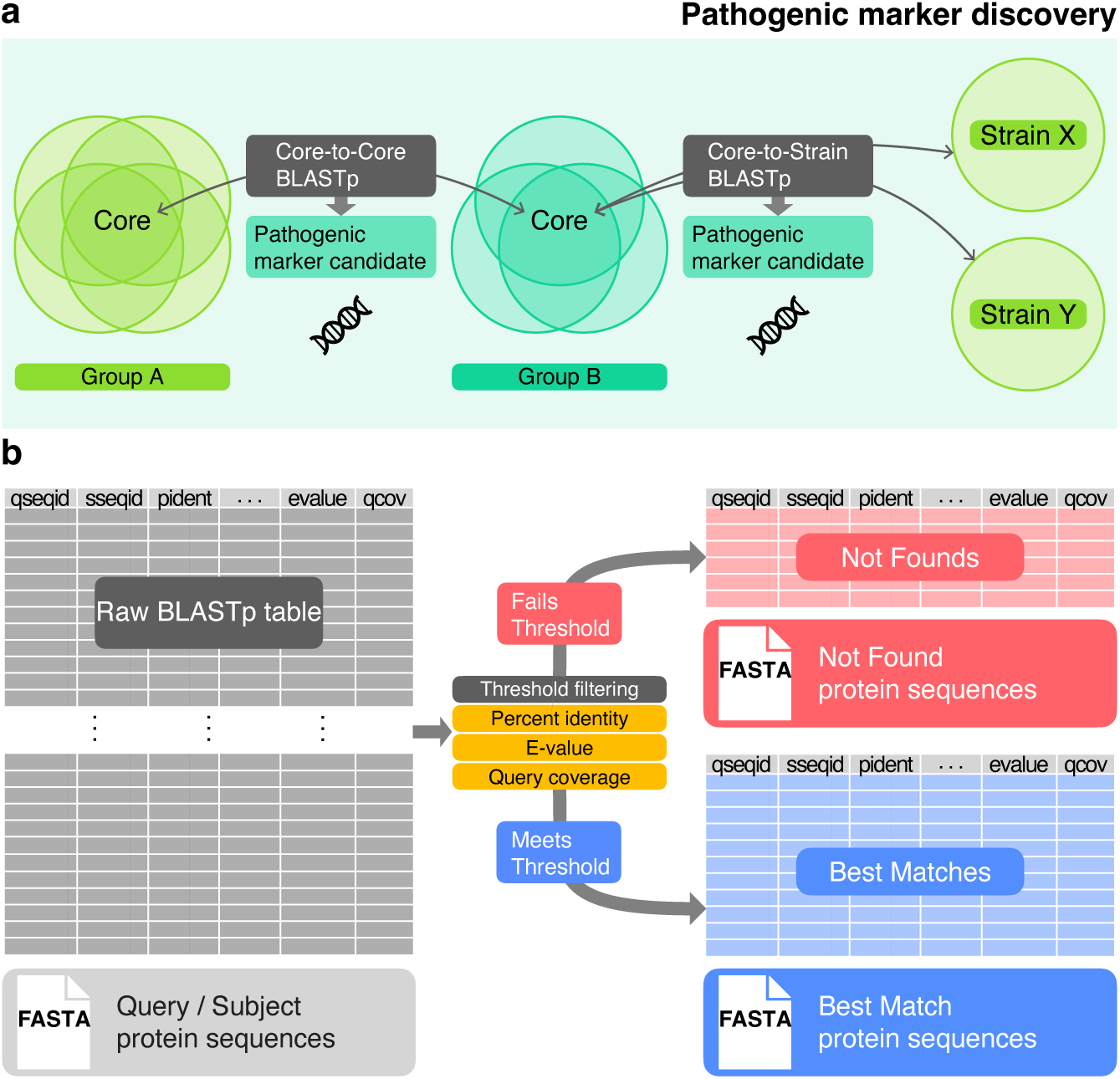
Pathogenic Marker Discovery module in ABComp. **a** Schematic of two comparison strategies supported by ABComp: In Core-to-Core analysis, the query group core genes are aligned to subject group core genes via BLASTp. In Core-to-Strain analysis, the query group core genes are aligned to all genes of each strain in the subject group via BLASTp. These strategies enable detecting of genes conserved in the query group but absent in the subject group. **b** Determination of gene presence and absence based on BLASTp alignment results. Genes that pass user-defined thresholds for percent identity, E-value, and query coverage are recorded as “Best Matches”, while genes that fail all alignments are classified as “Not Founds” and serve as group-specific marker candidates.

In the Assembly Polishing module (Fig. 1a), ABComp takes as input a draft long-read genome assembly generated by Flye^10^ or Unicycler^14^, as well as Illumina paired-end short reads. Trimmomatic^31^ is applied to short reads for adapter trimming, followed by quality control with FastQC (https://www.bioinformatics.babraham.ac.uk/projects/fastqc/) and MultiQC^32^ for summarized reports. Analysis-ready short reads are then employed for assembly polishing with Polypolish^33^, improving base-level accuracy of long-read draft assemblies, especially in repetitive or homopolymeric regions. The completeness and quality of the polished genome are evaluated using BUSCO^34^ and QUAST^35^, which provide standardized assembly metrics and reports. Notably, it is critical to optimize the initial long-read assembly beforehand to ensure maximum continuity and completeness of the polished genome sequences (Methods).

The Comparative Genomics module (Fig. 1b) operates on polished or pre-assembled genome sequences and initiates with gene annotation using Prokka^36^, a standardized bacterial annotation tool. This is followed by pairwise comparison of strains against a reference genome sequence using Roary^37^. In parallel, virulence and resistance factors are screened using ABRicate (https://github.com/tseemann/abricate) with a panel of curated databases^38–44^. Following strain-level analyses, the gene annotations are grouped according to predefined strain-to-group category information. Within each group, Roary performs pangenome analysis to identify core and shell gene sets. Functional annotation of non-hypothetical protein-coding genes is performed using EggNOG-mapper v2^45^, allowing classification into Cluster of Orthologous Genes^46,47^ (COG) categories and providing visual summaries of functional composition across groups. Together, we developed the ABComp pipeline to detect core virulence genes from bacterial genomes through automated assembly polishing and comparative genomics modules.

### Pathogenic marker discovery module of ABComp enables detection of core virulence factors from multiple bacterial genomes

To detect core virulence factors from pathogenic bacteria through phenotype-driven genomic analysis, we added the final module of ABComp for discovery of pathogenic markers from pathogens. The final module of ABComp enables flexible identification of pathogenic marker candidates by verifying gene presence and absence through pairwise protein sequence alignment using BLASTp^48^ (Fig. 2). Notably, unlike the preceding automated modules, this step is intentionally user-guided to allow custom comparisons tailored to specific biological questions or phenotype-driven groupings.

ABComp proposes two comparison strategies within the Pathogenic Marker Discovery module, leveraging the core and shell gene sets defined in the Comparative Genomics module. In the Core-to-Core analysis, the core genes of the query group are aligned against those of a subject group to identify genes that are highly conserved in the query group yet absent in the subject group (Fig. 2a). In the Core-to-Strain analysis, the query group’s core genes are aligned against the full gene annotation of each strain in the subject group. The collected results identify query group core genes that are consistently absent across all subject strains (Fig. 2a). To determine gene presence or absence, ABComp applies tunable thresholds based on percent identity, E-value, and query coverage of amino acid sequences for each gene. Genes meeting all criteria in at least one alignment are retained as ‘Best Matches’ with the top-scoring hit, while those failing the threshold for all alignments are labeled as ‘Not Found’ (Fig. 2b). These ‘Not Found’ genes are interpreted as absent in the subject group and serve as critical gene candidates for the query group. This framework enables hypothesis-driven marker discovery, supporting downstream functional analysis or experimental validation.

The modular structure allows ABComp to accommodate either raw sequencing data or pre-assembled genomes (Fig. 1b), enabling scalable and customizable analyses that bridge genome assembly, comparison, and phenotype-driven pathogenic marker discovery in a unified framework. Table 1 summarizes all bioinformatics tools and versions employed in ABComp. More detailed descriptions of each module process can be found in the Methods section. In summary, ABComp is an optimized tool for determining critical factors through phenotype-driven comparative genomic analysis.

### ABComp accurately detects yersiniabactin locus from *Klebsiella pneumoniae* ground truth dataset

To evaluate the functionality and performance of ABComp, we utilized a recently published, publicly available dataset of *Klebsiella pneumoniae*^49^. This dataset comprised sequencing reads generated from both short-read (Illumina) and long-read (ONT) platforms. Antibiotic-resistant *K. pneumoniae* is critical due to its significant global impact on nosocomial infections and associated mortality^6,50^. The virulence of *K. pneumoniae* is highly associated with mobile genetic elements, among which the integrative and conjugative element ICE*Kp* is the most prevalent^51^. ICE*Kp* carries key virulence loci, including the yersiniabactin (ybt) locus, a cluster of genes that encodes the proteins necessary for production of the siderophore yersiniabactin. This siderophore scavenges iron from the surroundings, enhancing the survival of the pathogen within the host and providing an advantage against host defense mechanisms^51,52^. Previously in the original study, the ybt locus was detected by Kleborate^53^, a tool for various screening analyses of *K. pneumoniae* genomes. We grouped the strains possessing the ybt locus as high-virulent (hv), while the others as low-virulent (lv). This classification served as a ground truth reference to evaluate whether ABComp could accurately detect ybt locus genes as hv-specific marker genes.

We first utilized ONT long reads to build draft assemblies using Flye or Unicycler from 42 deposited *K. pneumoniae* genomes. A two-step criterion was applied to the assembly results to assess the success of assembly optimization (Methods). Then, the successful draft assemblies and short reads were used as input for the Assembly Polishing module. Subsequently, the polished genome sequences were analyzed based on hv and lv groups in the Comparative Genomics module. We achieved high-quality genomes, confirmed by BUSCO and QUAST analyses, with minimal fragmented or missing genes (Fig. 3a) and nearly complete and continuous assemblies (Supplementary Table 1,2). Pangenome analysis using Roary revealed distinct core and shell gene sets for each group (Supplementary Fig. 1). Functional annotation of core and shell genes using EggNOG-mapper v2 revealed enrichment in metabolism, cellular processing, and transport among hv-core genes, alongside a proportion of hypothetical proteins (Fig. 3b and Supplementary Fig. 2a,b). The core and shell genes of the lv group also showed similar COG proportions to those of the hv group (Fig. 3b and Supplementary Fig. 2c,d). Interestingly, although the proportions of COG categories in both hv and lv groups were similar, nearly 1,000 more genes were identified as shell genes within the lv group (Fig. 3b). This observation aligns with the phenomenon where bacteria undergoing genome reduction often enhance their pathogenicity by eliminating superfluous genes, thereby streamlining their genomes for more specialized, virulent functions^54^. Finally, ABRicate results using the Virulence Factor Database (VFDB) further validated the presence of ybt locus genes within all hv strains, yet they were absent from all strains in the lv group (Supplementary Fig. 3).

**Figure 3.**
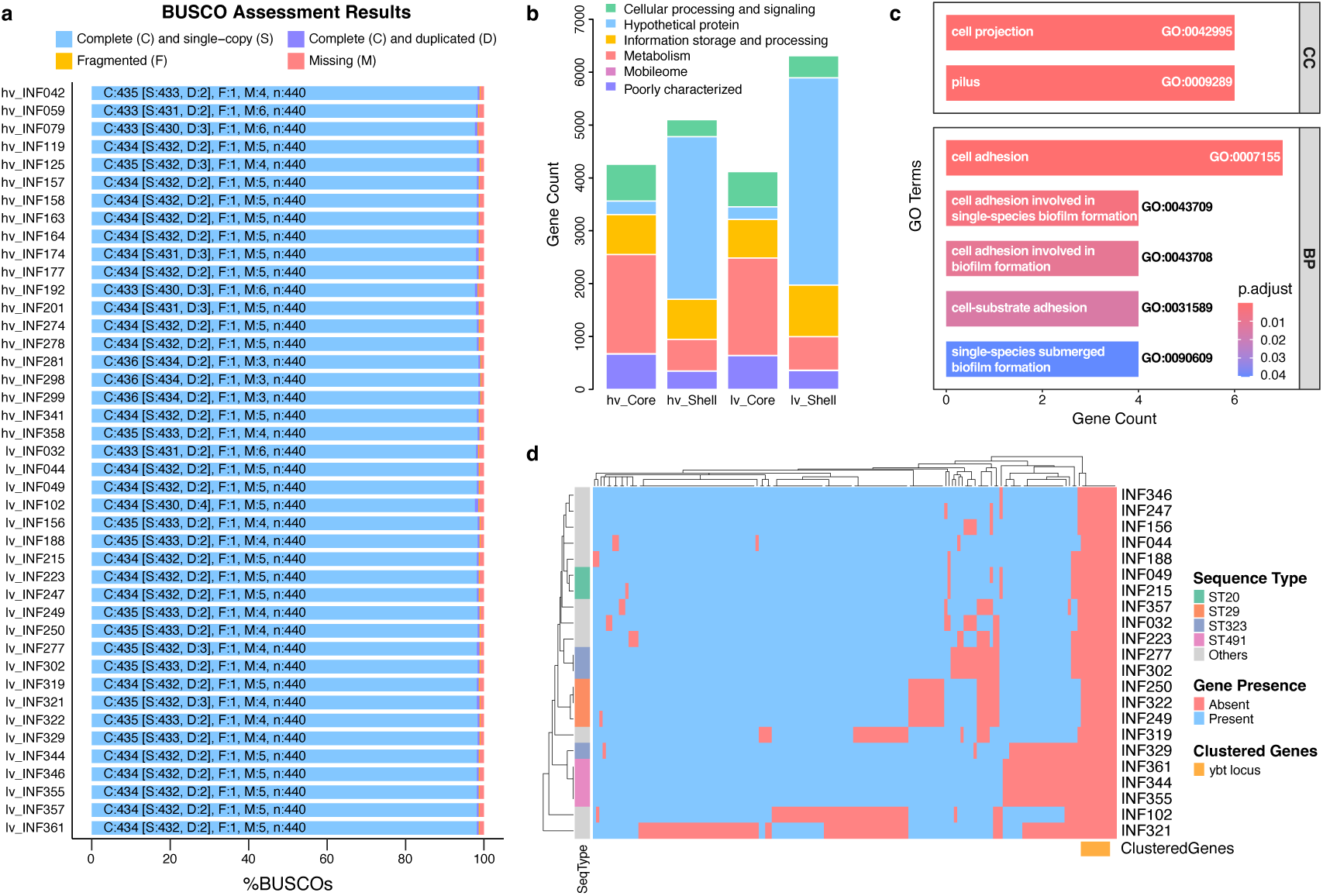
ABComp detects ybt locus and virulence-associated traits from *Klebsiella pneumoniae* strains. **a** BUSCO genome assessment plot. Most strains show >95% completeness and minimal fragmented or missing genes. **b** COG proportion of hv and lv core and shell genes, broadly classified into 5 functional supergroups. **c** Gene Ontology overrepresentation analysis (GO-ORA) of hv-core-specific genes reveals significant enrichment in virulence-related pathways such as cell adhesion, biofilm formation, and pilus assembly. **d** Heatmap showing presence or absence of hv-core genes across lv strains. Genes absent from all lv strains include the ybt locus genes.

To identify potential group-specific pathogenic genes, the hv and lv groups were further compared using the Pathogenic Marker Discovery module. First, the Core-to-Core analysis was conducted by setting the hv group as the query (hv-core) and the lv group as the subject group (lv-core). A total of 123 hv-core genes were identified as “Not Found” within the lv-core set (Supplementary Table 3). Therefore, these represent the hv-core-specific genes. We further questioned the functional relevance of these hv-core-specific genes and conducted Gene Ontology^55^ overrepresentation analysis (GO-ORA) using the complete hv-core gene set as background (Methods). Notably, cell adhesion and biofilm formation were significantly enriched biological processes among the hv-specific genes, and cell projection and pilus cell components were also identified (Fig. 3c). These pathways are well-known virulence traits in *K. pneumoniae* that enhance bacterial colonization, immune evasion, and persistence within host tissues^56^. Thus, genes associated with cell adhesion and biofilm formation pathways may exhibit stronger correlation within the hv group, suggesting their role as stabilized functional traits essential for enhanced virulence. This finding demonstrates that ABComp not only correctly identifies the ybt locus as a ground-truth marker but also uncovers functionally relevant hv-specific pathways associated with increased virulence potential in clinical bacterial isolates. Further Core-to-Strain analysis revealed the strain-wise distribution of hv-core genes. Importantly, nine annotated genes within the ybt locus, including well-known virulence factors such as *ybtS*, *ybtU*, and *fyuA*, were completely absent in all 22 lv strains, confirming their hv-specificity and validating ABComp’s accuracy in detecting known virulence factors (Fig. 3d). Collectively, these results demonstrate that ABComp successfully identifies both well-established virulence markers and pathogenic pathways, providing comprehensive insights into group-specific virulence mechanisms in clinical isolated pathogens.

### ABComp unbiasedly uncovers key virulence determinants in hypervirulent *K. pneumoniae*

To demonstrate ABComp’s ability to identify phenotype-relevant pathogenic markers in an untargeted manner, we analyzed 39 pre-assembled *K. pneumoniae* clinical isolate genomes with experimentally measured LD_50_ values from a mouse pneumonia model^57^. The study defined convergent strains featuring both multi-drug resistance and virulence factors found in hypervirulent *K. pneumoniae* (hvKP) strains^57^. Despite their genomic characteristics including mucoid receptors or siderophores, convergent strains showed unexpectedly low virulence in the mouse model^57^ (Fig. 4a). Given this, we hypothesized that hvKP strains possess other conserved pathogenic markers contributing to low LD_50_ values which ABComp could identify.

**Figure 4.**
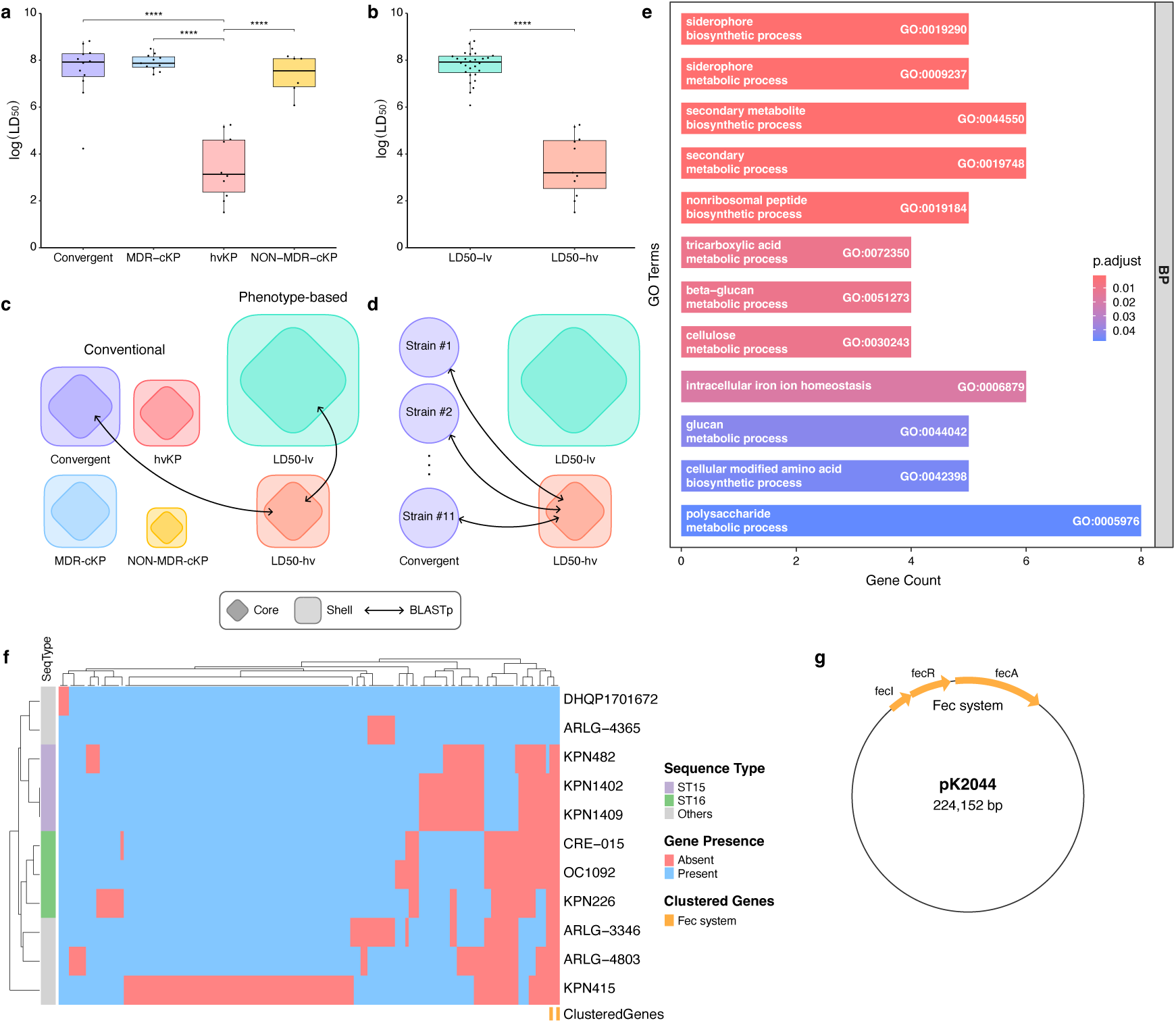
ABComp highlights the ferric citrate uptake system as a conserved pathogenicity marker in low-LD_50_ *K. pneumoniae* strains. **a** LD_50_ values of 39 *K. pneumoniae* strains classified by conventional groups (Convergent, hvKp, MDR-cKP, Non-MDR-cKP), adapted from *Kochan et al., 2023*. Convergent strains show unexpectedly low virulence despite harboring canonical virulence loci. **b** Phenotypic grouping of the 39 strains into LD_50_-hv and LD_50_-lv groups based on LD_50_ values from an *in vivo* mouse infection assay. Each data point represents an individual isolate (*n* = 39). Boxes indicate interquartile ranges, center lines denote medians, and whiskers show minima and maxima. Data for each isolate represent the mean of at least two independent experiments, each with three technical replicates. Statistical analyses were performed using one-way ANOVA with Tukey’s multiple-comparison test for (**a**) and a two-tailed Student’s *t*-test for (**b**); *p*-values are indicated as \*\*\*\**p* ≤ 0.0001. **c** Schematic overview of two Core-to-Core analyses utilizing the conventional group setting and the phenotype-based group setting. **d** Core-to-Strain analysis between the LD_50_-hv-core gene set and Convergent strains. **e** Gene Ontology overrepresentation analysis (GO-ORA) of 140 LD_50_-hv-specific core genes, compared to LD_50_-lv core genes. Bars are colored by adjusted *p*-values. **f** Presence and absence of LD_50_-hv-core genes across 11 Convergent strains. Hierarchical clustering was performed using Euclidean distance and complete linkage. **g** Genetic organization of the three genes within Fec system located on plasmid pK2044 (224,152 bp).

To test this hypothesis, we executed the ABComp comparative genomics module using two group settings: conventional groups (Convergent, hvKP, MDR-cKP (classical KP), Non-MDR-cKP) from the literature^57^, and custom phenotype-based groups (LD_50_-hv for low LD_50_/high virulence and LD_50_-lv for high LD_50_/low virulence) (Fig. 4a,b) (Methods). Using the ABComp pathogenic marker discovery module, Core-to-Core analysis between LD_50_-hv and LD_50_-lv groups revealed 140 LD_50_-hv-specific core genes (Fig. 4c) (Supplementary Table 4). GO-ORA of LD_50_-hv-specific core genes showed significant enrichment in siderophore metabolism and extracellular polysaccharide metabolism pathways (Fig. 4e), consistent with the hypermucoviscosity characteristic of LD_50_-hv strains^56,58,59^.

We then compared the Convergent group of the conventional setting and the LD_50_-hv group of our custom setting. There was only one strain, K014, which had an overlap between the two groups. Core-to-Core analysis between the LD_50_-hv and Convergent group revealed 179 LD_50_-hv-specific genes (Supplementary Table 5) and GO-ORA showed enrichment in siderophore and amino acid metabolism (Fig. 4c, Supplementary Fig. 4). Core-to-Strain analysis using LD_50_-hv core genes against Convergent strain annotations (excluding K014) revealed three contiguous plasmid-encoded genes absent in 9 out of 11 Convergent strains (Fig. 4d,f). These genes comprised the ferric citrate uptake (Fec) system, including sigma factor FecI and receptor gene FecA (Fig. 4f,g). The sigma factor regulator FecR was also absent in the same 9 strains but was not within the LD_50_-hv core set. The Fec system is an iron uptake pathway distinct from the siderophore-mediated systems, providing iron scavenging capacity under iron-limiting conditions such as host environments^60,61^. Because the Fec system enhances bacterial survival and competitiveness under host-associated iron limitation, and has also been linked to increased fitness and virulence in multidrug-resistant *K. pneumoniae* strains^62^, we selected the Fec system as a potential contributing factor for increased virulence in the LD_50_-hv group and conducted experimental validation. Together, these results demonstrate ABComp’s ability to uncover gene candidates associated with high virulence in an untargeted manner.

## Discussions

Bacterial pathogens continue to evolve under selective pressures from host immunity, antimicrobial exposure, and environmental adaptations, giving rise to both multidrug resistant and hypervirulent lineages. While whole-genome sequencing technology has transformed the ability to capture this bacterial genetic diversity, translating genomic data into clinically actionable insights remains challenging, especially when the goal is to identify genomic markers that distinguish one clinical phenotype from another.

ABComp is a flexible and automated pipeline dedicated its functionality for phenotype-driven bacterial group comparisons. ABComp unites hybrid assembly polishing, pangenome-based comparative genomics within user-defined groups, and pathogenic marker discovery into a single workflow. The pipeline is modularized to accept both raw sequencing reads, as well as pre-assembled genome sequences. As public repositories continue to accumulate vast amounts of diverse bacterial sequencing data, ABComp’s automated and group-level comparison framework will enable researchers to widely leverage this growing resource for precise, phenotype-focused discoveries.

Applied to a curated ground truth dataset of *Klebsiella pneumoniae* clinical isolates grouped by the presence and absence of the yersiniabactin (ybt) locus, ABComp successfully identified the entire locus as high-virulence group-specific core set. Functional enrichment analysis of the high-virulence group-specific core genes highlighted biological processes such as cell adhesion and biofilm formation, suggesting that the conservation of these traits is correlated to the presence of siderophores, and may contribute to enhanced pathogenic potential.

The ability to move beyond known virulence factors and pinpointing phenotype-driven pathogenic markers is a major strength of ABComp. By applying the pipeline to clinical isolates with experimentally measured virulence, we identified the plasmid-encoded ferric citrate (Fec) system as a potential hypervirulence-associated marker candidate. Such findings exemplify how ABComp can detect both established and novel group-specific markers, offering new hypothesis for functional validation and potential targets for therapeutic interventions.

## Methods

### Assembly polishing module

The assembly polishing module of ABComp requires a draft assembly built from long reads (e.g. Oxford Nanopore Technologies, Pacific Biosciences), and short reads (e.g. Illumina paired-end reads) to polish the draft assembly. For building the draft assembly from long reads, either Flye^10^ or Unicycler^14^ is recommended. Building a draft assembly often requires a manual parameter tuning process to validate the completeness of the assembly, especially the continuity. For example, tuning the minimum read overlap parameter in Flye affects the continuity of the genome, due to the read length distribution of long reads. Therefore, this preliminary assembly tuning process is omitted from the automation, so the user can input the optimized draft assembly result from manual parameter tuning. The assembly polishing module starts with assessing the quality of raw paired-end short reads with FastQC (https://www.bioinformatics.babraham.ac.uk/projects/fastqc/) (v0.12.1) and adapter trimming with Trimmomatic^31^ (v0.39). The trimmed reads are once again assessed through FastQC. The FastQC reports for all short reads, both raw and trimmed, are each merged through MultiQC^32^ (v1.24.1), resulting in a single quality report for all input reads. Next, adapter-trimmed short reads are employed to polish the draft assembly with Polypolish^33^ (v0.5.0). Polypolish improves base-level accuracy in regions where long-read assemblies tend to be less precise, particularly in homopolymeric or repeat regions. After polishing, the quality of the polished genome is assessed, with BUSCO^34^ (v5.5.0) and QUAST^35^ (v.5.2.0). BUSCO uses lineage datasets to evaluate the completeness of genome assemblies by comparing the presence of certain key genes in the assembly. QUAST reports metrics to evaluate the completeness of the genome assembly.

### Comparative genomics module

The Comparative Genomics module operates from complete, analysis-ready genomes. The genomes can be the output of the Assembly Polishing module, or the user can provide pre-assembled genomes to analyze. The genome and corresponding GenBank file of a reference strain is required, to aid the gene annotation process and conduct a pairwise comparison analysis with all input strains. The overall gene content of the genomes is annotated using Prokka^36^ (v1.14.6). The resulting gene feature files (GFF) from Prokka are directly compared to the reference genome annotation, using Roary^37^ (v3.12.0). This process eases the matching of genes between the reference strain and the individual strains. Then, the GFFs are copied and grouped into each group directory, making use of the group and strain information initially provided by the configuration file. Within each group directory, Roary conducts a pangenome analysis, using the previously grouped gene feature files. This process identifies the core and shell genes of each group, based on identity parameters the user set beforehand. In this work, for simplicity, we define ‘core’ genes as the union of core and soft-core genes identified by Roary, representing genes present in default setting of 95% or more of all strains within a group. Similarly, we define ‘shell’ genes as the union of shell and cloud genes of Roary, representing genes present in less than default setting of 95% of all strains within a group. For each group, the protein FASTA files are summarized for core and shell gene protein products. During this procedure, ABComp employs a systematic approach to deal with hypothetical proteins. Hypothetical proteins are proteins inferred through the open reading frames (ORFs) found in the assemblies, but no known function or similar protein of annotation has been identified in existing databases^63^. Therefore, the ratio of hypothetical proteins within the total protein-coding annotation can serve as an indicator of how well the strain’s functional profile is understood. To deal with hypothetical proteins, ABComp first generates three types of core and shell FASTA files: with all genes, without hypothetical proteins, and only hypothetical proteins (6 combinations). The functional annotation is then proceeded to the core and shell protein sets without the hypothetical proteins. EggNOG-mapper v2^45^ (v2.1.12) is used to query these proteins to pre-downloaded EggNOG databases, resulting in functional annotation including Clusters of Orthologous Groups^46,47^ (COG), Gene Ontology^55^ (GO), and Kyoto Encyclopedia of Genes and Genomes^64^ (KEGG) pathways. ABComp specifically focuses on COG analysis, because it helps categorize proteins by evolutionary lineage, which is essential for understanding bacterial functional diversity and adaptation. First, a pie plot is generated representing the overall ratio of the 26 COG categories, and these categories are broadly grouped into five functional supergroups. During this process, the held-out hypothetical proteins are integrated into the pie plot, thereby visualizing the ratio of hypothetical proteins altogether with the COG ratio of the remaining core, shell protein sets (Supplementary Fig. 2). These results provide a functional overview of each group’s core and shell genes. Parallel from functional annotation, the annotation results from Prokka are also screened for the presence of antimicrobial genes or virulence factors using ABRicate (v1.0.1). In ABComp, we dealt with preselected databases that are compatible with ABRicate: NCBI AMRFinderPlus^38^, ARG-ANNOT^39^, CARD^40^, Megares^41^, Resfinder^42^, PlasmidFinder^43^ and VFDB^44^. The query results of these databases are summarized in each input strain directory. This process highlights the content of clinically relevant genes that may not be captured in standard gene ontology or COG analysis, thus enhancing the overall understanding of each bacterial strain while ensuring accurate detection of resistance genes.

### Pathogenic marker discovery

The Pathogenic Marker Discovery module requires a query group and a subject group for both Core-to-Core and Core-to-Strain analysis. In both analyses, the subject group set is made into a database using the makeblastdb command of BLASTp^48^ (v2.15.0). Next, the core gene set of the query group is queried to the database with specifying the query coverage option. This produces a raw BLASTp table showing the total alignment and statistics of the query entry. ABComp requires the thresholds for three main metrics, percent identity, E-value and query coverage. Users can set the pass or fail hyperparameters to define the presence or absence of the query entry within the subject database. In this work, a query was considered present if at least one BLASTp alignment satisfied all the following criteria: >35% identity, E-value < 0.001, and >80% query coverage. On the other hand, queries in which all the alignments fail the thresholds were considered absent. We termed these queries as “Not Founds”.

### Grouping strains

For the *K. pneumoniae* ground truth dataset, we grouped the 42 strains into high-virulence (hv) and low-virulence (lv) groups based on the presence of the ybt locus. The hv group had 20 strains, with various types of ybt locus originating from either ICEKp (*n* = 17) or plasmids (*n* = 3). The presence of the ybt locus, within ICE*Kp* or plasmids, was hypothesized to be accurately detected by ABComp as a hv-specific core gene. The lv group had 22 strains, lacking any type of ybt locus. In this setting, we did not differentiate between the various types of ybt loci or ICE*Kp* subtypes. The analysis was solely focused on the presence of the ybt locus as the defining criterion for grouping strains into hv and lv categories.

For the *K. pneumoniae* LD_50_ phenotype dataset, we first utilized the conventional grouping from the original paper, which is based on genomic features and antibiotic resistance; Convergent (*n* = 12), hvKP (*n* = 10), MDR-cKP (*n* = 11), and NON-MDR-cKP (*n* = 6). For the custom phenotypic classification, we classified the 39 strains into LD_50_-hv (*n* = 11) and LD_50_-lv (*n* = 28) groups by K-means clustering of LD_50_ values, setting K to 2.

### Generating draft assembly

Long reads were initially assembled using Flye (v2.8.1) adjusting the min-overlap parameter, with default set to 5000. Successful assemblies were defined by two-step criteria. The initial criterion: a single circular contig (∼5 Mbp) identified as the chromosome, with additional circular contigs considered plasmids, as well as the absence of fragmented contigs or repeats. Of the 44 strains, 35 met the initial criterion and were confirmed successful. For the nine strains failing the initial criterion, a secondary criterion was applied: when a single circular contig (∼5Mbp) was identified with fragmented contigs, but with no repeats, fragmented contigs smaller than 10 kbp were discarded. 5 strains passed the second criteria and were additionally confirmed successful. Finally, the remaining four strains were assembled with Unicycler (v0.4.8), utilizing the hybrid assembly option. Before running Unicycler, the short reads were trimmed with Trimmomatic (v0.39) for removal of adapter sequences. The two-step criteria mentioned for Flye were equally applied to Unicycler results. Among the four strains, two strains passed the criteria and were confirmed successful. The final two strains resulted in short fragmented contigs with repeats for both Flye and Unicycler, were excluded. Finally, Medaka (https://github.com/nanoporetech/medaka) (v1.11.3) was used with the default model option to polish the 42 strains. The parameter value for Flye and Unicycler, as well as number of contigs and their circularity for each strain can be found at Supplementary Table 1.

### Gene Ontology Overrepresentation Analysis

To investigate the functional implications of genes that are present in one group but absent in the other, we performed Gene Ontology overrepresentation analysis (GO-ORA). Direct GO term assignments for background genes were obtained using FaCoP.v2^65^, a functional annotation pipeline that integrates diverse biological knowledgebases to infer Gene Ontology terms. Among the initial background genes, we filtered genes that were mapped to at least one GO term as the background gene set. The direct GO annotation was further expanded using the buildGOmap() function for inferring indirect annotation. GO-ORA was conducted with the gene set of interest using the clusterProfiler^66^ (v4.0.5) R package, with Biological Process (BP) and Cellular Component (CC) ontologies. The enricher() function was used to identify significantly enriched GO terms, with Benjamini-Hochberg correction applied for multiple testing. Terms with adjusted *p*-value < 0.05 were considered statistically significant.

## Data availability

The ONT and Illumina raw reads of *K. pneumoniae* ybt ground truth dataset can be found at the European Nucleotide Archive (ENA), under BioProject accessions PRJEB6891 and PRJNA351909. The pre-assembled genomes for *K. pneumoniae* LD_50_ phenotype dataset is uploaded at GenBank, under BioProject accessions <u>PRJNA788509</u>.

## Code availability

ABComp and the codes used in this study, is publicly available on https://github.com/young5454/ABComp.git

## Acknowledgments

We are grateful to all of authors and Yeom lab members for comments on the early draft of manuscript. This work was supported by the National Research Foundation Korea Basic Science Research Programs (2021R1C1C1005184, 2020M3A9H5104237 and 2020R1A5A1019023 to J.Y.), Creative-Pioneering Researchers Program through Seoul National University, the Korea Health Technology R&D Project through the Korea Health Industry Development Institute (KHIDI) (grant number: HI23C026400 and RS-2023-00304637), the Global-LAMP Program of the National Research Foundation of Korea (NRF) grant funded by the Ministry of Education (No. RS-2023-00301976).

