## Supplementary Figures for "A Snakemake-based bacterial whole genome comparison pipeline for multi-group clinical isolates"

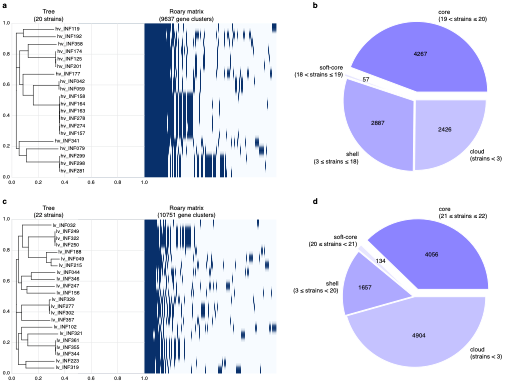

**Supplementary Figure 1. Group-wise pangenome analysis of 42 *K. pneumoniae* isolates in ybt ground truth dataset with Roary. a** Pangenome matrix for the hv group (20 strains) with an accompanying phylogeny tree. Each column is a gene cluster (total 9,637 clusters), dark blue indicates presence. **b** Pangenome composition of the hv group. Segment labels indicate gene counts. **c** Pangenome matrix for the lv group (22 strains)(total 9,637 clusters). **d** Pangenome composition of the lv group.

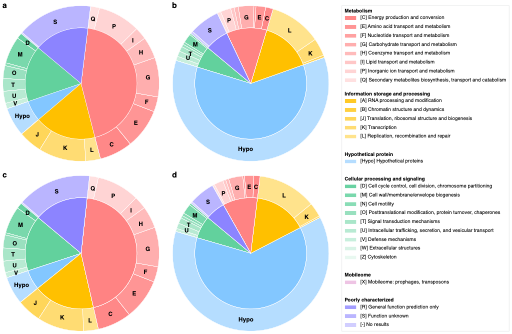

**Supplementary Figure 2. Group-wise COG functional annotation of 42 *K. pneumoniae* isolates in ybt ground truth dataset with EggNOG-mapper v2. a** hv core genes. **b** hv shell genes. **c** lv core genes. **d** lv shell genes. Inner rings show the five COG supergroups, outer rings show the 26 COG categories plus hypothetical proteins (denoted as “Hypo”). Categories contributing <1% of total genes are unlabeled for clarity. The legend lists full COG names and letter codes.

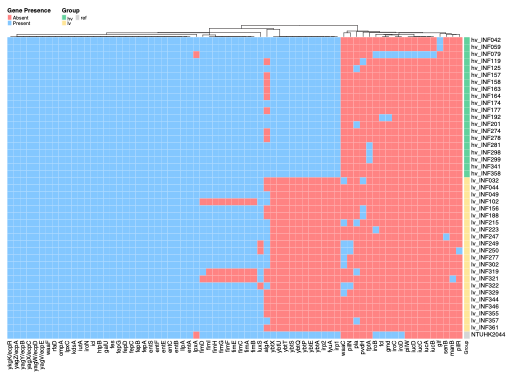

**Supplementary Figure 3. Virulence factor screening of 42 *K. pneumoniae* isolates in ybt ground truth dataset with ABRicate.** Heatmap showing presence or absence of virulence factors using the Virulence Factor Database (VFDB). Virulence factors are clustered according to hierarchical clustering of presence/absence profiles. The hv and lv groups are clearly distinguished based on ybt locus genes (*ybtX*, *ybtU*, *ybtT*, *ybtS*, *ybtQ*, *ybtP*, *ybtE*, *ybtA*, *irp2*, *fyuA*).

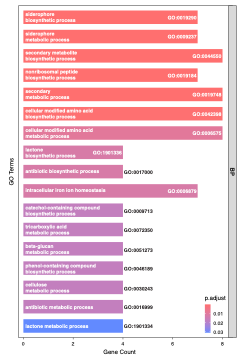

**Supplementary Figure 4. Gene Ontology overrepresentation analysis (GO-ORA) of 179 LD_50_-hv-specific core genes, compared to Convergent core genes.** Bars are colored by adjusted *p*-values.
